# A Test of the Pioneer Factor Hypothesis

**DOI:** 10.1101/2021.08.17.456650

**Authors:** Jeffrey L Hansen, Barak A Cohen

## Abstract

The Pioneer Factor Hypothesis (PFH) states that pioneer factors (PFs) are a subclass of transcription factors (TFs) that bind to and open inaccessible sites and then recruit non-pioneer factors (nonPFs) that activate batteries of silent genes. We tested the PFH by expressing the endodermal PF FoxA1 and nonPF Hnf4a in K562 lymphoblast cells. While co-expression of FoxA1 and Hnf4a activated a burst of endoderm-specific gene expression, we found no evidence for functional distinction between these two TFs. When expressed independently, both TFs bound and opened inaccessible sites, activated endodermal genes, and “pioneered” for each other, although FoxA1 required fewer copies of its motif to bind at inaccessible sites. A subset of targets required both TFs, but the mode of action at these targets did not conform to the sequential activity predicted by the PFH. From these results we propose an alternative to the PFH where “pioneer activity” depends not on the existence of discrete TF subclasses, but on TF binding affinity and genomic context.

## Main

Transcription factors (TFs) face steric hindrance when instances of their motifs are occluded by nucleosomes ^1,2^. This barrier prevents spurious transcription but must be overcome during development when TFs activate batteries of silent genes. The PFH describes how TFs recognize and activate nucleosome-occluded targets. According to the PFH, specialized subclasses of TFs collaborate sequentially to activate their targets. Pioneer factors (PFs) bind to and open inaccessible sites and then recruit non-pioneer factors (nonPFs) that are responsible for recruiting additional factors to initiate gene expression ^3–6^.

PFs also play a primary role in cellular reprogramming by first engaging silent regulatory sites of ectopic lineages ^7^. Continuous overexpression of PFs and nonPFs can lead to a variety of lineage conversions ^8–13^. The conversion from embryonic fibroblasts to induced endoderm progenitors offers one clear example ^12,13^. This reprogramming cocktail combines the canonical PF FoxA1 ^6^ and nonPF Hnf4a ^14^ and is suggested to rely upon sequential PF and nonPF behavior ^15^. We used this cocktail to test the PFH.

The PFH makes strong predictions about the activities of ectopically expressed PFs and nonPFs. Because PFs are defined by their ability to bind nucleosome-occluded instances of their motifs, the PFH predicts that PFs should bind to a large fraction of their motifs. However, similar to other TFs, PFs only bind a limited subset of their inaccessible motifs ^16–19^. There are chromatin states that are prohibitive to PF binding ^17,20^ and, in at least two cases, FoxA1 requires other TFs to bind its sites ^18,21^. These examples suggest that PFs are not always sufficient to open inaccessible chromatin. The PFH also predicts that nonPFs should only bind at accessible sites, yet the bacterial protein LexA can pioneer inaccessible sites in mammalian cells ^22^. These observations, and the absence of direct genome-wide interrogations of the PFH, prompted us to design experiments to test major predictions made by the PFH using FoxA1 and Hnf4a as a model PF and nonPF.

To test these predictions, we expressed FoxA1 and Hnf4a separately and together in K562 lymphoblast cells and then measured their effects on DNA-binding, chromatin accessibility, and gene activation. In contrast to the predictions of the PFH, we found that both FoxA1 and Hnf4a could independently bind to inaccessible instances of their motifs, induce chromatin accessibility, and activate endoderm-specific gene expression. The only notable distinction between the two factors was that Hnf4a required more copies of its motif to bind at inaccessible sites. When expressed together, co-binding could only be explained in a minority of cases by sequential FoxA1 and Hnf4a activity. Instead, most co-bound sites required concurrent co-expression of both factors, which suggests cooperativity between these TFs at certain repressive genomic locations. We propose an alternative to the PFH that eliminates the distinction between PFs and nonPFs and instead posits that the energy required to pioneer occluded sites (“pioneer activity”) comes from the cumulative affinity of motifs and cooperativity between TFs.

## Results

### Generation of FoxA1 and Hnf4a clonal lines

We tested predictions of the PFH using FoxA1 as a model endoderm PF and Hnf4a as a model nonPF. Because PFs are defined by their behavior in ectopic settings, we expressed FoxA1 and Hnf4a in mesoderm derived K562 lymphoblast cells. These cells express neither FoxA1 nor Hnf4a and present an entirely new complement of chromatin and co-factors. Thus any ectopic signature that we observe is due primarily to the TFs themselves. We focused only on the initial response to TF expression to capture primary mechanisms of TF behavior and not the secondary effects that can lead to cellular conversion and that may confound our analyses.

To perform these experiments, we created lentiviruses that inducibly express either FoxA1 or Hnf4a (Fig. 1A). We created cassettes in which a doxycycline inducible promoter drives either FoxA1 or Hnf4a and cloned these cassettes separately into a lentiviral vector ^23^ that constitutively expresses Green Fluorescent Protein (GFP). Although PFs are typically expressed at supraphysiological levels ^24,25^, we infected K562 cells with each vector at a multiplicity of infection (MOI) of one to limit the degree of non-specific effects. We then used flow cytometry to sort single cells and selected FoxA1 and Hnf4a clones that had similar GFP levels to ensure that our clones carried a similar transgene load. Finally, we performed both doxycycline titration induction and time course experiments to identify the minimum doxycycline concentration and treatment time for robust TF activity. We observed that 0.5 µg/ml doxycycline for 24 hours was the minimal treatment condition that allowed *FoxA1* and *Hnf4a*, and their respective target genes *ALB* and *APOB*, to reach a plateau of expression (Supplementary Fig. 1). We used these conditions in our subsequent experiments.

**Fig. 1:**
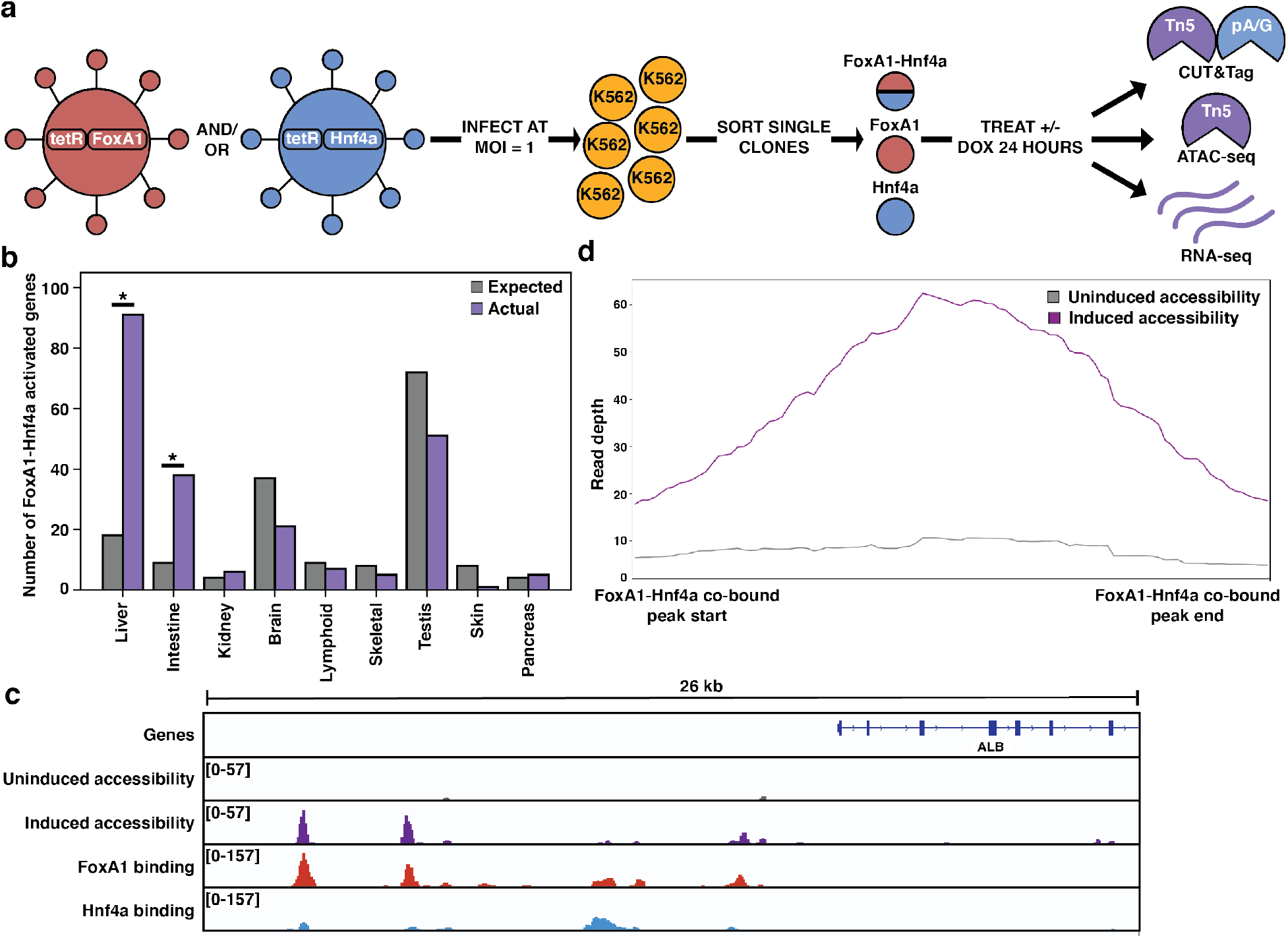
FoxA1-Hnf4a pioneers liver-specific loci in K562 cells. **(a)** Schematic of experimental design to infect K562 cells with FoxA1- or Hnf4a-lentivirus and then perform functional assays on dox-induced cells. In CUT&Tag, a protein A-protein G fusion (pA/G) increases the binding spectrum for Fc-binding and allows Tn5 recruitment to antibody-labeled TF binding sites. In ATAC-seq, Tn5 homes to any accessible site. And in RNA-seq, polyA RNA is captured and sequenced. **(b)** The number of tissue-specific genes predicted from the hypergeometric distribution to be activated by FoxA1-Hnf4a compared to the number actually activated. Both liver-(*P* < 10^−38^) and intestinal-enrichment (*P* < 10^−13^) are significant. There are 242 total liver-enriched genes and 122 total intestine-enriched genes. **(c)** Genome browser view of a representative liver-specific locus (*ALB*) in FoxA1-Hnf4a clonal line that shows uninduced and induced accessibility, FoxA1 binding, and Hnf4a binding. **(d)** Meta plot showing uninduced and induced accessibility at all FoxA1-Hnf4a co-bound sites within 50 kb of each FoxA1-Hnf4a activated liver-specific gene (n = 53).

### Co-expression of FoxA1 and Hnf4a in K562 cells conforms to the predictions of the PFH

The first prediction of the PFH is that co-expression of FoxA1 and Hnf4a should be sufficient to induce ectopic tissue-specific gene expression. We tested this prediction by infecting our FoxA1 clonal line with Hnf4a-expressing lentivirus to generate a double expression clonal line, hereafter referred to as FoxA1-Hnf4a. Upon co-induction in K562 cells we observed strong enrichment for both liver- and intestine-specific gene activation; FoxA1-Hnf4a activated 91 liver-specific genes (18 expected, *P* < 10^−38^, cumulative hypergeometric) and 38 intestinal genes (9 expected by chance, *P* < 10^−13^, cumulative hypergeometric) (Fig. 1B). The dual liver and intestine enrichment that we observed is consistent with the finding that intestinal gene regulatory networks appear during reprogramming experiments that aim to use FoxA1-Hnf4a to convert embryonic fibroblasts to the liver lineage ^13^. We conclude that FoxA1 and Hnf4a are sufficient to activate endoderm-specific gene expression in the ectopic K562 line.

Where ectopic genes are activated in K562 cells, the PFH predicts co-binding of FoxA1 and Hnf4a at inaccessible sites and induction of chromatin accessibility. Alternatively, FoxA1 and Hnf4a may not be able to overcome the K562 chromatin environment and instead activate gene expression by binding exclusively to accessible K562 sites. To distinguish between these possibilities, we measured FoxA1 and Hnf4a binding by CUT&Tag ^26^ after induction, and chromatin accessibility by ATAC-seq ^27^ both before and after doxycycline induction. At the liver-specific locus *Albumin* (*ALB)*, FoxA1 and Hnf4a co-bound at inaccessible sites and increased accessibility (Fig. 1C). This pattern was consistent surrounding FoxA1-Hnf4a activated liver genes: 43 of the 53 co-bound sites within 50 kb of a FoxA1-Hnf4a activated gene were inaccessible prior to induction, and the average accessibility signal at these co-bound sites increased substantially upon induction (Fig. 1D).

Although we focused on functional binding surrounding activated liver genes, these patterns were consistent across the genome. The vast majority of both FoxA1 and Hnf4a binding sites fell within inaccessible regions (Supplementary Fig. 2) and both FoxA1 and Hnf4a opened the majority of the inaccessible sites to which they bound (Supplementary Fig. 2). These results show that despite an entirely ectopic complement of chromatin and co-factors within mesoderm derived K562 cells, the endodermal TFs FoxA1 and Hnf4a can find and activate the correct genes. Most individual binding by FoxA1 and Hnf4a near their co-activated genes occurred at the same sites bound in HepG2 liver cells ^28^ (Supplementary Fig. 2). Altogether we conclude that when co-expressed, FoxA1 and Hnf4a conform to the predictions of the PFH and that cis-regulatory sequences are sufficient to guide their activity within an ectopic cell type.

### Both FoxA1 and Hnf4a individually activate many liver-specific genes

We next sought to test whether ectopic tissue-specific gene expression in K562 cells results from the sequential activity of FoxA1 and Hnf4a as predicted by the PFH. Sequential activity assumes that Hnf4a won’t bind and FoxA1 won’t activate, therefore neither FoxA1 nor Hnf4a should activate tissue-specific gene expression when expressed alone. To test this prediction, we induced K562 lines expressing either FoxA1 or Hnf4a alone and measured mRNA expression by RNA-seq. FoxA1 induction resulted in strong liver-specific enrichment (*P* < 10^−4^, cumulative Hypergeometric) and weak intestinal-specific enrichment (not significant) (Fig. 2A), while Hnf4a induction resulted in both strong liver-specific enrichment (*P* < 10^−8^, cumulative Hypergeometric) and strong intestinal-specific enrichment (*P* < 10^−15^, cumulative Hypergeometric) (Fig 2B). Importantly, neither FoxA1 nor Hnf4a are expressed within K562 cells nor did they induce expression of the other TF, suggesting that the expression changes we observed were due to the independent effects of either FoxA1 or Hnf4a.

**Fig. 2:**
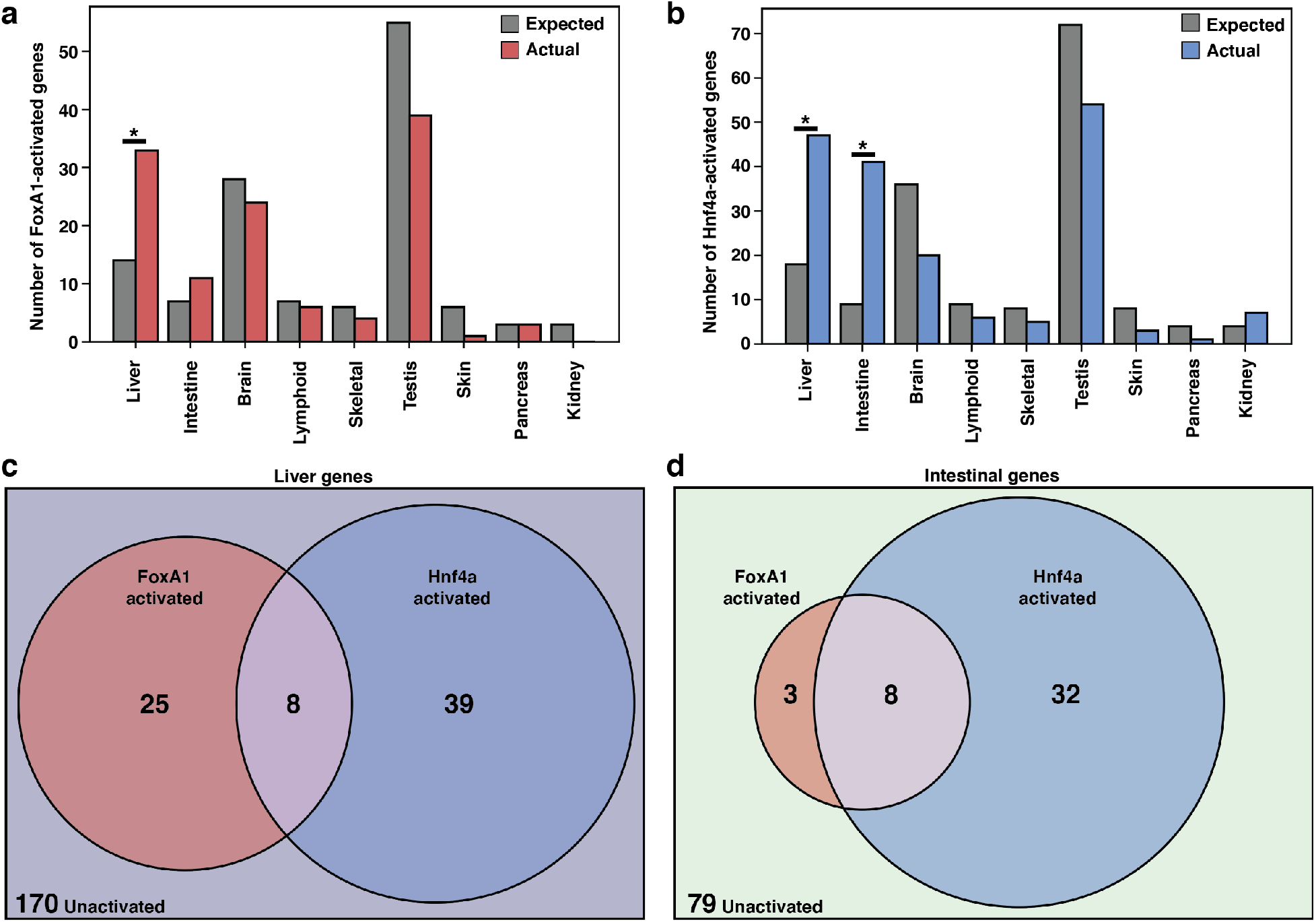
FoxA1 and Hnf4a activate independent liver- and intestine-specific genes. **(a)** The number of tissue-specific genes predicted from the hypergeometric distribution to be activated by FoxA1 compared to the number actually activated. Liver-enrichment (*P* < 10^−4^) is significant. There are 242 total liver-enriched genes. **(b)** The number of tissue-specific genes predicted from the hypergeometric distribution to be activated by Hnf4a compared to the number actually activated. Liver-(*P* < 10^−8^) and intestine-enrichment (*P* < 10^−15^) are significant. There are 242 total liver-enriched genes and 122 total intestine-enriched genes. **(c)** 242 liver genes characterized as activated by Foxa1, Hnf4a, both, or neither. **(d)** 122 intestine genes characterized as activated by FoxA1, Hnf4a, both, or neither.

When expressed individually, FoxA1 and Hnf4a activated largely independent sets of liver genes (Fig. 2C) and intestinal genes (Fig. 2D). FoxA1 activates liver genes enriched for fibrinolysis and complement activation (Supplementary Table 1) whereas Hnf4a activates liver genes enriched for cholesterol import and lipoprotein remodeling (Supplementary Table 2). Thus, in contrast to the predictions of the PFH, FoxA1 and Hnf4a are each sufficient to induce separate and specific endodermal responses when expressed alone in K562 cells.

### Both FoxA1 and Hnf4a can independently bind and open inaccessible sites around liver genes

Our results raised the possibility that both FoxA1 and Hnf4a can pioneer inaccessible instances of their motifs. To test this possibility, we induced FoxA1 and Hnf4a expression individually and then measured each factor’s binding profile and their accessibility profiles before and after induction. FoxA1 induction resulted in FoxA1 binding and induced accessibility adjacent to *Arg1*, a liver-specific gene that is silent in K562 cells (Fig. 3A), while Hnf4a alone bound and induced accessibility at sites nearby the liver-specific gene *ApoC3* (Fig. 3B). This pattern was consistent across liver-specific loci. 34 of the 59 FoxA1 binding sites within 50 kb of a FoxA1-activated liver gene were inaccessible and opened upon induction (Fig. 3C) as was the case for 39 of the 76 Hnf4a binding sites (Fig. 3D). We observed similar patterns genome-wide. FoxA1 and Hnf4a bound primarily to inaccessible sites (Supplementary Fig. 3), opened them (Supplementary Fig. 3), and in regions surrounding activated genes, most binding occurred at the same sites bound in HepG2 liver cells (Supplementary Fig. 3). We conclude that FoxA1 and Hnf4a have roughly equivalent abilities to bind and open inaccessible sites.

**Fig. 3:**
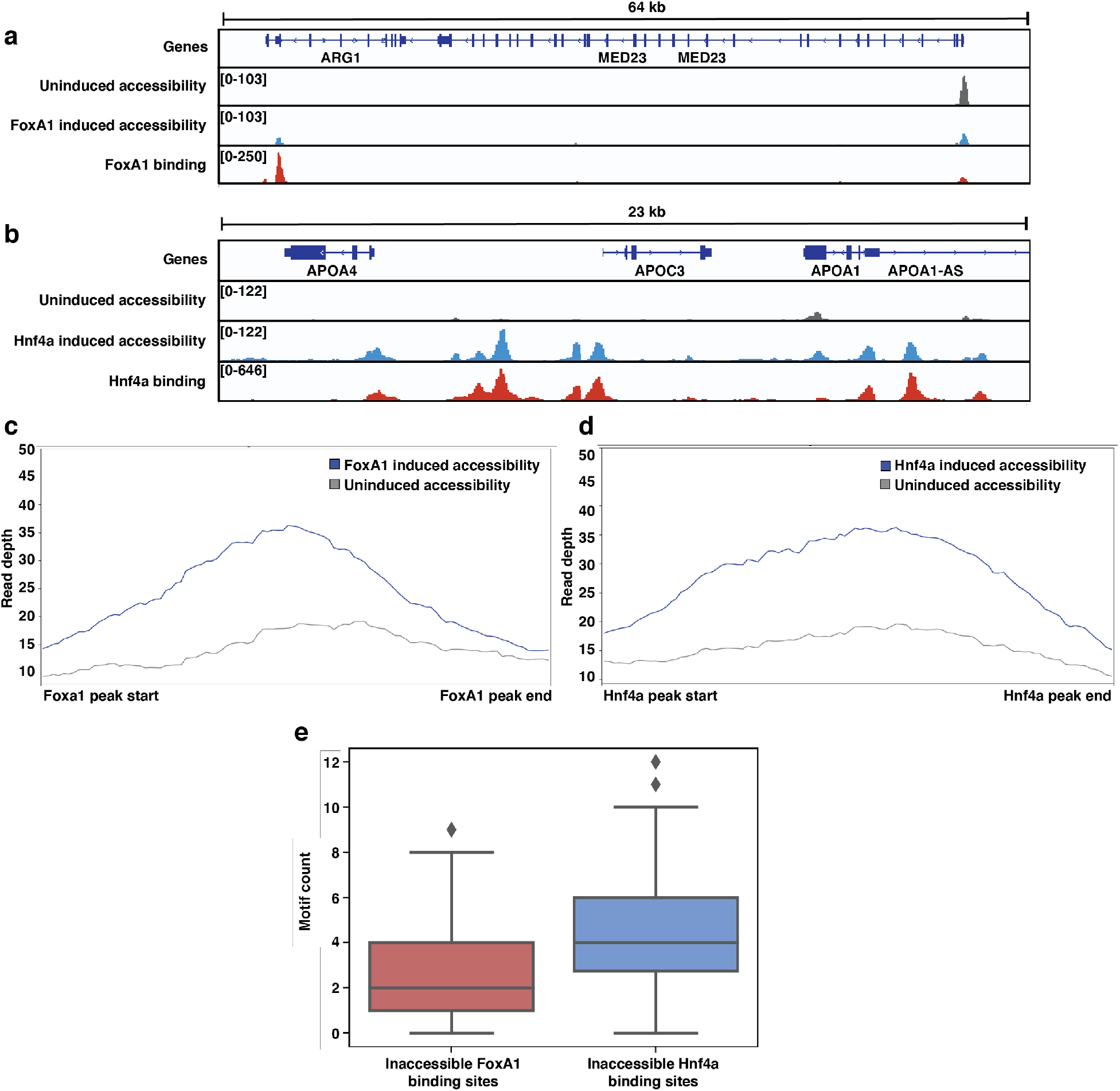
Both FoxA1 and Hnf4a can pioneer liver-specific loci. **(a)** Genome browser view of a representative liver-specific locus (*Arg1*) in FoxA1 clonal line showing uninduced and induced accessibility and FoxA1 binding. **(b)** Genome browser view of a representative liver-specific locus (*ApoC3*) in Hnf4a clonal line showing uninduced and induced accessibility and Hnf4a binding. **(c)** Meta plot of uninduced and induced accessibility at all FoxA1 binding sites within 50 kb of each FoxA1-activated liver-specific genes (n = 59). **(d)** Meta plot of uninduced and induced accessibility at all Hnf4a binding sites within 50 kb of each Hnf4a-activated liver-specific genes (n = 76). **(e)** FoxA1 or Hnf4a motif count at FoxA1 or Hnf4a binding sites within 50 kb of each FoxA1- or Hnf4a-activated liver-specific genes, respectively. Motifs were called with FIMO using 1e-3 a p-value threshold. For each boxplot, the center line represents the median, the box represents the first to third quartiles, and the whiskers represent any points within 1.5 times the interquartile range.

Because this finding was incompatible with the current formulation of the PFH, we sought to understand how we might reconsider the factors’ behavior. We used FIMO (MEME Suite) ^29^ with JASPAR motif matrices (Supplementary Fig. 4) ^30^ to examine the motif content at sites bound by either FoxA1 or Hnf4a in K562 cells. Sites where FoxA1 and Hnf4a showed independent pioneering activity contained occurrences of each factor’s cognate motif. Sites independently pioneered by FoxA1 contained between 1-4 motifs, while sites pioneered by Hnf4a contained 3-6 motifs (Fig. 3E). This is despite the fact that the FoxA1 motif occurs more frequently across the genome than the Hnf4a motif (Supplementary Fig. 4). This observation is consistent with data that show that FoxA1 binds with stronger affinity than Hnf4a ^31–33^ and suggests that “pioneer activity” may depend on the cis-regulatory context and not on special subclasses of TFs.

### Some liver genes require collaborative FoxA1-Hnf4a activity

In addition to those genes independently activated by Foxa1 and Hnf4a, there is an additional set of 31 liver genes that are not activated until both FoxA1 and Hnf4a are present (Fig. 4A). We therefore asked whether the activation of these 31 liver genes conforms to the PFH. If these genes conform to the PFH, then we would expect each target to have nearby sites where FoxA1 binds individually and where FoxA1 and Hnf4a co-bind when expressed together. We have called these sites “FoxA1 Pioneered” (FP). Sites are “Hnf4a Pioneered” (HP) if Hnf4a binds individually and FoxA1 and Hnf4a co-bind when expressed together and sites are “Collaboratively Co-bound” (CC) if neither TF binds individually but both do when expressed together. There are examples of each modality surrounding AMDHD1, a liver-specific gene co-activated by FoxA1 and Hnf4a (Fig. 4B). When we examine all of the liver genes only activated by FoxA1-Hnf4a co-expression, we find that in contradiction with the PFH, there are roughly equal numbers of FP, HP, and CC sites (Fig. 4C). Therefore, in most cases, genes that require joint FoxA1-Hnf4a activity do not rely on FoxA1 pioneer activity.

**Fig. 4:**
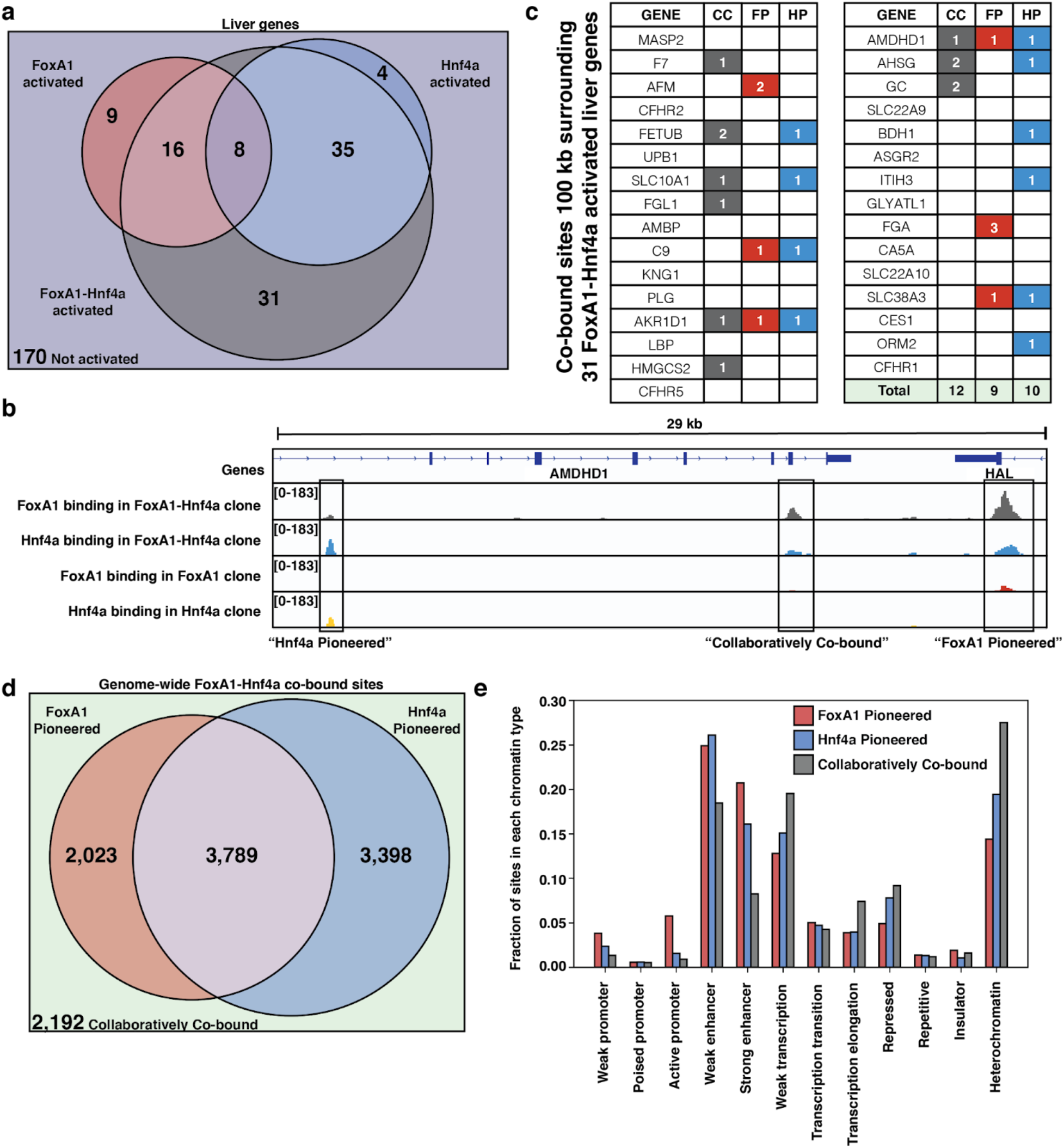
FoxA1 and Hnf4a both pioneer and collaborate at liver-specific sites. **(a)** Venn diagram of all liver genes categorized as either activated by FoxA1, Hnf4a, FoxA1-Hnf4a, some combination, or by none of the three cocktails. **(b)** Genome browser view of a representative liver-specific locus (*AMDHD1*) showing examples of a co-bound site that is “FoxA1 Pioneered” (FP), “Hnf4a Pioneered” (HP), and “Collaboratively Co-bound” (CC). The first two tracks are FoxA1 and Hnf4a binding in the FoxA1-Hnf4a co-expression clone and the last two tracks are FoxA1 and Hnf4a binding in their individual expression clones. **(c)** List of the 31 liver genes that are only activated by FoxA1-Hnf4a co-expression. The columns indicate how many co-bound FP, HP, or CC peaks exist within 100 kb of the gene. **(d)** Venn diagram of all genome-wide co-bound peaks categorized as either bound by FoxA1 individually (FP), Hnf4a individually (HP), by both, or by neither (CC). **(e)** Overlap of FP, HP, and CC sites from (D) with ChromHMM annotations showing the fraction of each co-binding site type in each chromatin region.

The patterns of genome-wide co-binding and accessibility of FoxA1 and Hnf4a follow similar trends. Of the 11,402 co-bound sites, 2,023 were FP, 3,398 were HP, and 2,192 were CC (Figure 4D) and FoxA1-induced differentially accessible peaks explain a minority of the FoxA1-Hnf4a differentially accessible peaks (Supplementary Fig. 5). Collaborative co-binding may be necessary in less accessible parts of the region, as there are more CC sites in ChromHMM-labeled ^34^ heterochromatic and repressed regions, and there are more FP and HP sites in promoter and enhancer regions (Fig. 4E).

## Discussion

In contrast to the predictions of the PFH, we found that both canonical PF FoxA1 and nonPF Hnf4a can independently bind nucleosome-occluded sites, increase accessibility, and activate nearby endodermal genes. Other endodermal genes require the combined activity of both factors, but the mode of action at these targets does not conform to the predicted sequential activity of FoxA1 followed by Hnf4a. These observations suggest an alternative model to the PFH during endodermal reprogramming in which FoxA1 and Hnf4a each independently activate a unique set of genes, and also collaborate, perhaps through cooperative binding, at another distinct set of targets.

Our results support efforts to revisit the independent activities of TFs in reprogramming cocktails. Early reprogramming of fibroblasts to myoblasts relied solely upon the ectopic overexpression of MyoD ^25,35^ and new reprogramming cocktails have been tested and validated in a large-scale screen for single, cell autonomous reprogramming TFs ^24^. Increasing the efficiency of reprogramming cocktails that depend on multiple TFs will require distinguishing between the independent and cooperative effects of TFs. For example, our finding that Hnf4a independently activates more intestine-specific genes than FoxA1 raises the possibility that titrating down Hnf4a activity during reprogramming could result in a more liver-specific profile. Such fine-tuning of TF activities has been suggested as an option to improve the success of other reprogramming cocktails ^36–38^.

While we did not find evidence for a clear distinction between the functional activities of FoxA1 and Hnf4a, our results do suggest that FoxA1 may require fewer copies of its motif than Hnf4a to elicit a response. This could be because FoxA1 has stronger affinity for its motif than Hnf4a. FoxA1 has a three-dimensional shape that is hypothesized to compete with histones ^39^ and the measured affinity of FoxA1 for its motif is higher than that of Hnf4a for its motif ^31–33^. Thus, FoxA1’s designation as a PF and Hnf4a as a nonPF may be due to FoxA1 having a stronger affinity for DNA than Hnf4a.

Although we found clear instances of sites independently pioneered by either FoxA1 or Hnf4a, not all sites containing multiple motifs were pioneered in K562 cells, which comports with studies showing that the sequence context in which motifs occur also plays an important role in determining whether sites will be pioneered or not. Gal4’s ability to bind nucleosomal DNA templates depends both on the number of copies of its motif ^40^ and the positioning of the motif in the nucleosome ^41^. Precise nucleosome positioning also dictates TP53 and Oct4 pioneering behavior ^42,43^. A TF’s motif affinity, motif count, and the presence of co-factor motifs are all strong predictors of pioneer activity ^18,19,44–48^ and certain types of heterochromatic patterning have been labeled “pioneer resistant” ^17^. Pioneer activity may best be summarized then by the free energy balance between TFs, nucleosomes, and DNA ^49,50^ rather than as a property of specific classes of TFs.

## Methods

### Cloning, production, and infection of viral vectors

We used PCR to add V5 epitope tags to the 3’ end of FoxA1 (Addgene #120438) and Hnf4a (Addgene #120450) constructs and then used HiFi DNA Assembly (NEB #E2621L) to clone each construct into a pINDUCER21 doxycycline-inducible lentiviral vector (Addgene #46948). All primers are listed in Table 1. The Hope Center Viral Vector Core at Washington University in St. Louis then generated and titered high-concentration virus. We infected human K562 cells at a multiplicity of infection of 1 by spinoculation at 800G for 30 minutes in the presence of 10 µg/ml polybrene, passaged the cells for 3 days, and then selected for positively infected cells by single cell sorting on GFP+ into 96-well plates. Finally we used qPCR to select for clones that had high inducibility of TF and target gene expression (Supplementary Fig. 1).

### Cell culture

We grew K562 cells (ATCC CCL-243) in Iscove’s Modified Dulbecco Serum supplemented with 10% fetal bovine serum, 1% penicillin-streptomycin and 1% non-essential amino acids. When it was time to conduct one of our functional assays, we split FoxA1-, Hnf4a-, or FoxA1-Hnf4a-expressing cells into replicate flasks and then treated with +/- 0.5 µg/ml doxycycline for 24 hours.

### RNA extractions, reverse transcription, and qPCR

We extracted RNA from 1e6 cells/sample with the PureLink RNA Mini (Invitrogen #12183020) column extraction kit and completed on-column DNA digestion with PureLink DNase (Invitrogen #12185010). We quantified and assessed the quality of the RNA with an Agilent 2200 Tapestation instrument and then either froze down pure RNA for later RNA-sequencing library preparation or used ReadyScript cDNA Synthesis Mix (Sigma #RDRT-100RXN) to produce cDNA for qPCR. We performed qPCR with SYBR Green PCR Master Mix (Applied Biosystems #4301955) and gene-specific and housekeeping primers (Table 1).

### RNA-sequencing and analysis

We generated three replicates of +/- doxycycline-treated RNA-sequencing libraries with the NEBNext Ultra II Directional RNA Library Prep Kit (NEB #E7765S). We quantified and assessed the quality of the libraries with an Agilent 2200 Tapestation instrument, size selected with AMPure XP beads (Beckman Coulter #A63880), and then sequenced the libraries with 75bp paired-end reads on an Illumina NextSeq 500 instrument.

We quantified transcripts with Salmon ^51^, filtered out any with fewer than 10 reads, and then called differentially expressed transcripts with DeSeq2 ^52^. A gene was called differentially upregulated if it had a log2fold change of at least 1 and was called “activated” if it had fewer than 50 normalized reads in the uninduced control. A gene was called “tissue-specific” according to the Human Protein Atlas definition of tissue enrichment ^53^, which is if a gene is at least 4-fold higher expressed in the tissue-of-interest than in any other tissue.

### ATAC-sequencing and analysis

We followed the OMNI-Atac protocol ^54^ to generate two replicates of +/- doxycycline-treated low-background ATAC-sequencing libraries. We isolated 2e5 cells/sample and then extracted 5e4 nuclei/sample for tagmentation and library preparation. We quantified and assessed the quality of the libraries with an Agilent 2200 Tapestation instrument, size selected with AMPure XP beads, and then sequenced the libraries with 75bp paired-end reads on an Illumina NextSeq 500 instrument.

We aligned transcripts with bowtie2 ^55^ with the parameters: --local -X2000, generated RPKM normalized BigWig files for visualization with DeepTools bamCoverage ^56^, and then called peaks at low stringency with macs2 (p = 0.01) ^57^. With these peaks, we either called reproducible peaks with IDR (FDR of 0.05) ^58^ or used DiffBind ^59^ to call differential peaks.

### CUT&Tag and analysis

We followed the CUTANA Direct-to-PCR CUT&Tag protocol (EpiCypher) to generate two replicates of low-background CUT&Tag libraries. We isolated 1e5 cells/sample, and then either used rabbit anti-human FoxA1 monoclonal antibody (Cell Signaling #53528), mouse anti-human Hnf4a monoclonal antibody (Invitrogen #MA1-199), or rabbit anti-human histone H3K4me3 polyclonal antibody (Epicypher #13-0041) as a positive control. We amplified this signal with either goat anti-rabbit (Epicypher #13-0047) or goat anti-mouse (Epicypher #13-0048) polyclonal secondary antibodies. For a negative control, we omitted the primary antibody and checked for any non-specific pull-down. Finally, we used CUTANA pAG-Tn5 (Epicypher #15-1017) to tagment the genomic regions surrounding each bound antibody complex. We quantified and assessed the quality of the libraries with an Agilent 2200 Tapestation instrument, size selected with AMPure XP beads, and then sequenced the libraries with 150bp paired-end reads on an Illumina NextSeq 500 instrument.

When we assessed our libraries with the Agilent Tapestation instrument, we found that our negative controls had minimal signal. This is expected in the protocol and as such sequencing the sample is recommended as optional ^60^. For this reason, we sequenced only our positive samples. We aligned our samples with Bowtie2 ^55^ using recommended parameters ^60^: --very-sensitive --end-to-end --no-mixed --no-discordant -I 10 -X700, created RPKM normalized BigWig files with DeepTools bamCoverage ^56^, and called peaks with macs2 (p = 1e-5) ^57^ with recommended parameters ^26^. We then combined overlapping peaks from replicate samples using BEDTools intersect ^61^. We attributed binding sites to genes if they were within 50 kb (25 kb up- and 25 kb downstream) of the gene’s TSS. Because co-binding occurred less frequently, we attributed co-binding sites to genes if they were within 100 kb of the gene’s TSS. “FoxA1 Pioneered” sites were those where we identified overlapping FoxA1 and Hnf4a binding peaks within 100 kb of a gene that was only activated by FoxA1 and Hnf4a and where there was also an overlapping FoxA1 binding peak, when FoxA1 was expressed alone. “Hnf4a Pioneered” sites were those where we identified overlapping FoxA1 and Hnf4a binding peaks within 100 kb of a gene that was only activated by FoxA1 and Hnf4a and where was also an overlapping Hnf4a binding peak, when Hnf4a was expressed alone. And “Collaboratively Co-bound” sites were those where we identified overlapping FoxA1 and Hnf4a binding peaks within 100 kb of a gene that was only activated by FoxA1 and Hnf4a and where there was neither a FoxA1 nor Hnf4a binding peak.

### Tissue- and biological process-specific expression analysis

We generated lists of tissue-specific genes for each tissue by extracting “enriched genes” from the Human Protein Atlas. A tissue’s enriched genes are those whose mRNA expression is at least four-fold higher than expression found in any other tissue. We then computed hypergeometric assays to determine if our activated genes were enriched in any tissue-specific gene set. Finally, we used Panther gene ontology analysis to identify enriched biological processes.

### Genome tracks and profile plot analysis

We visualized the signal from our functional assays by loading each file into the Integrated Genome Viewer ^62^, using hg19 as reference. We then used the computeMatrix function in reference-point mode and plotProfile function, both with default parameters, in the DeepTools suite ^56^ to display aggregated CUT&Tag and ATAC-sequencing signals across indicated genomic regions.

### Motif and chromatin segmentation analysis

We used FIMO from the MEME Suite to identify occurrences of motifs. We used 1e-3 as a p-value threshold and JASPAR PWMs for FoxA1 (MA0148.1) and Hnf4a (MA0114.2). We used ChromHMM annotations ^34^ to characterize the epigenetic profile of FoxA1 and Hnf4a binding sites.

## Data Availability

All genomic sequencing data have been deposited on Gene Expression Omnibus (GEO) under accession number GSE182191.

## Acknowledgements

We thank Dr. Gary Stormo, Dr. Robi Mitra, and members of the Cohen Lab for reading and critiquing the manuscript and for helpful discussion; Jessica Hoisington-Lopez and MariaLynn Crosby in the DNA Sequencing Innovation Lab for assistance with high-throughput sequencing; the Genome Engineering and iPSC Center for allowing us to use their Sony Flow Cytometer for cell sorting; and Mingjie Li in the Hope Center Viral Vectors Core for assistance with producing lentiviral expression vectors. This work was supported by grants from the National Institutes of Health: R01GM092910 (Dr. Barak Cohen), T32HG000045 (Dr. Michael Brent, Washington University in St. Louis Genome Analysis Training Program), and T32GM007200 (Dr. Wayne Yokoyama, Washington University in St. Louis Medical Scientist Training Program).

## Author Contributions

J.L.H. and B.A.C. designed the overall project. J.L.H. conducted all experiments and analysis. J.L.H. and B.A.C. wrote the manuscript.

