## Supplementary information for "A Test of the Pioneer Factor Hypothesis"

#### Supplementary Figures

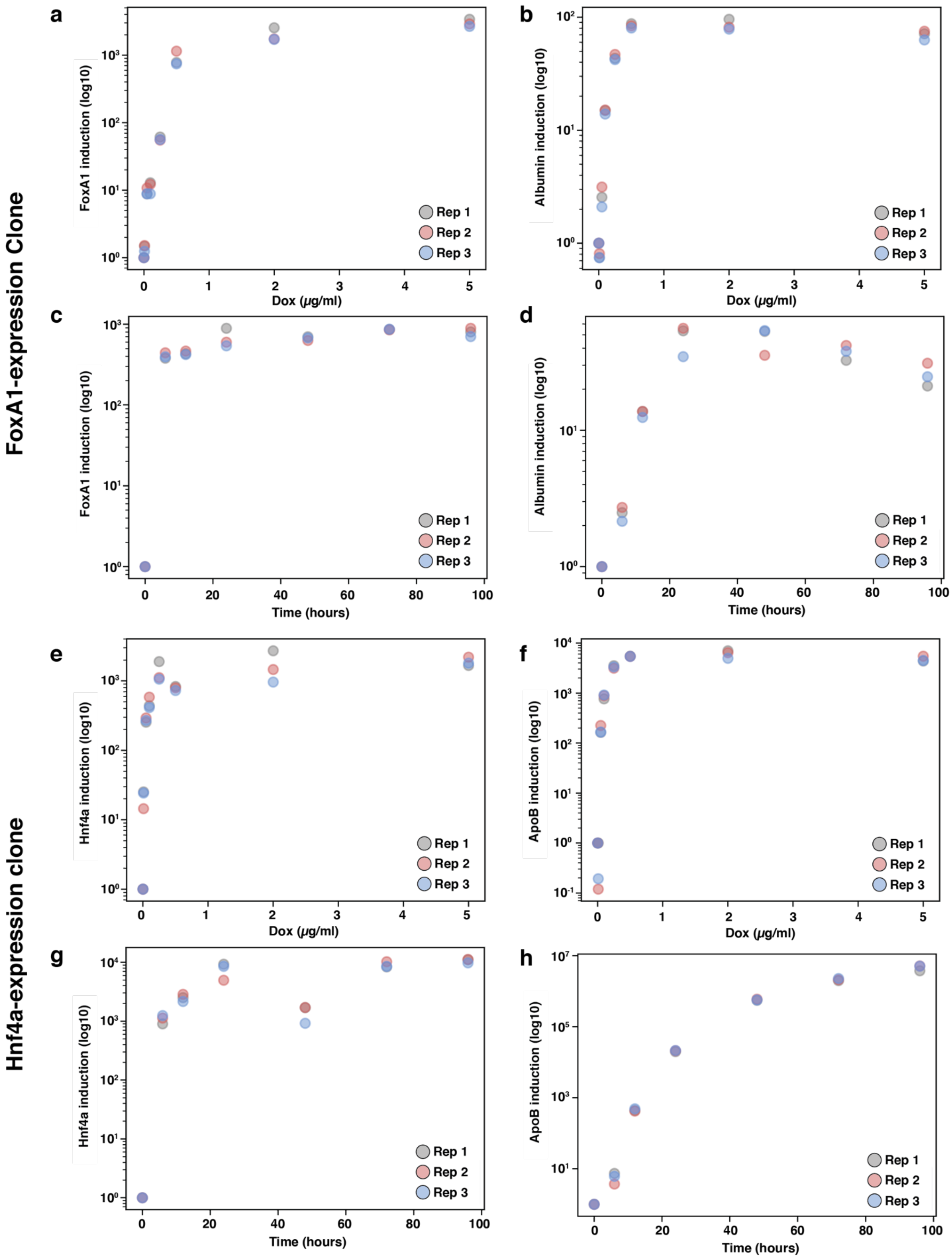
FoxA1-expression Clone
Hnf4a-expression clone
a
FoxA1 induction (log10)
Dox (µg/ml)
b
Albumin induction (log10)
Dox (µg/ml)
c
FoxA1 induction (log10)
Time (hours)
d
Albumin induction (log10)
Time (hours)
e
Hnf4a induction (log10)
Dox (µg/ml)
f
ApoB induction (log10)
Dox (µg/ml)
g
Hnf4a induction (log10)
Time (hours)
h
ApoB induction (log10)
Time (hours)
Rep 1
Rep 2
Rep 3

**Supplemental Fig. 1: Titration of doxycycline concentration and treatment time for TF and target gene induction.** qPCR measurements made from RNA extracted from either the FoxA1 clonal line **(a-d)** or the Hnf4a clonal line **(e-h)** that was treated with either increasing doxycycline concentrations or longer time periods. Expression is displayed as log10 fold induction over either 0 µg/ml doxycycline control (for concentration titration) or time 0 (for time titration). Each sample primer was normalized to the *HPRT* housekeeping gene. Doxycycline concentration titration measurements were made at 0, 0.01, 0.05, 0.1, 0.5, 2, and 5 µg/ml. Doxycycline treatment time measurements were made at 0, 6, 12, 24, 48, 72, and 96 hours.

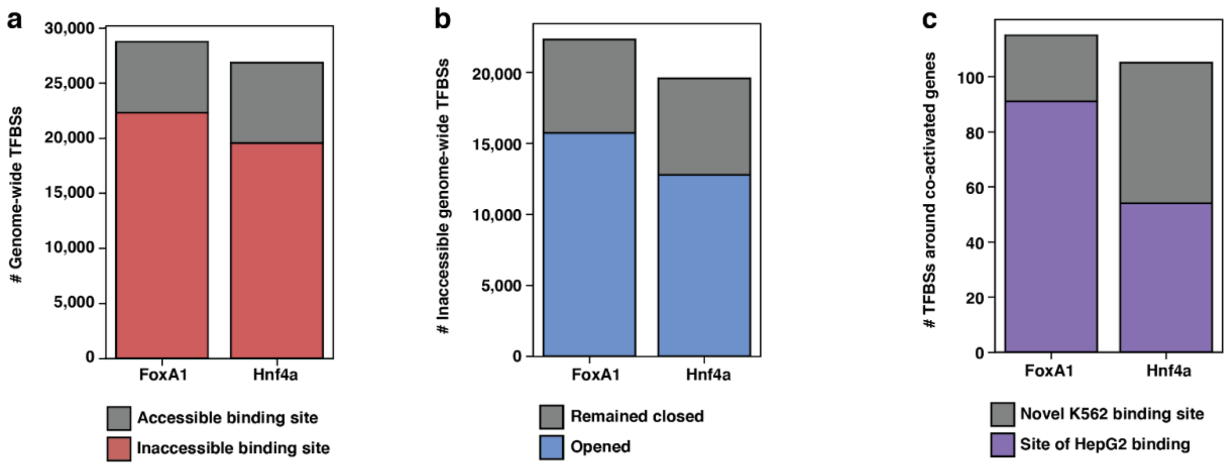
a
30,000
25,000
20,000
15,000
10,000
5,000
0
### Genome-wide TFBSs
FoxA1
Hnf4a
Accessible binding site
Inaccessible binding site
b
20,000
15,000
10,000
5,000
0
### Inaccessible genome-wide TFBSs
FoxA1
Hnf4a
Remained closed
Opened
c
100
80
60
40
20
0
### TFBSs around co-activated genes
FoxA1
Hnf4a
Novel K562 binding site
Site of HepG2 binding

**Supplemental Fig. 2: Characterization of FoxA1 and Hnf4a binding patterns in FoxA-Hnf4a clone.** **(a)** The number of genome-wide FoxA1 or Hnf4a TFBSs that overlap with an ATAC-seq peak in the uninduced (-dox) cells ("Accessible binding site") or that do not overlap with an ATAC-seq peak ("Inaccessible binding site"). **(b)** The number of inaccessible binding sites from (A) that overlap with an ATAC-seq peak in the induced (+dox) cells ("Opened") or that do not overlap with an ATAC-seq peak ("Remained closed"). **(c)** The number of FoxA1 or Hnf4a binding sites within 50 kb of each FoxA1-Hnf4a co-activated gene characterized as either a "HepG2 binding site," where the TFBS overlaps a TFBS of FoxA1 or Hnf4a in HepG2 liver cells, or as a "Novel K562 binding site," where the TFBS does not overlap with a HepG2 binding site.

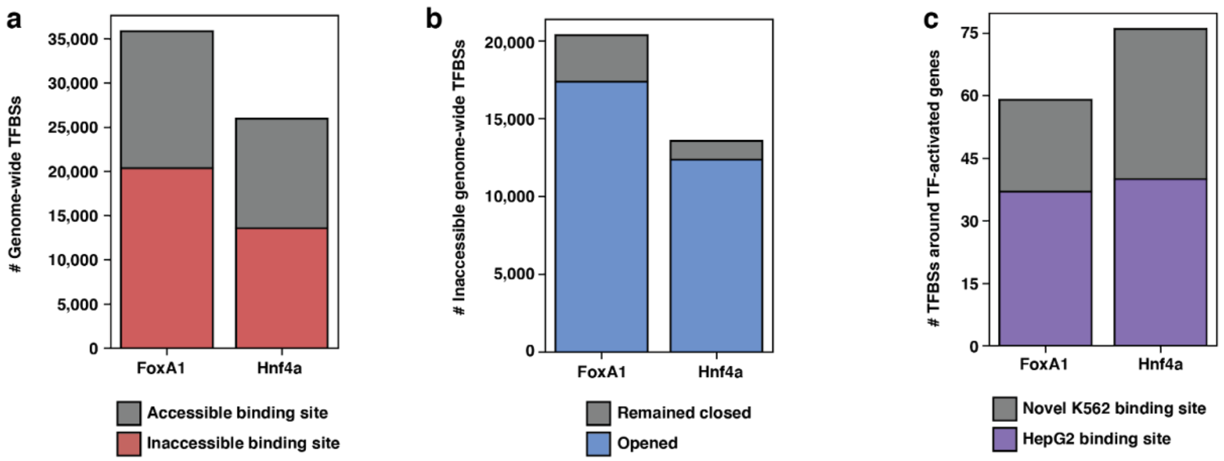
a
35,000
30,000
25,000
20,000
15,000
10,000
5,000
0
### Genome-wide TFBSs
FoxA1
Hnf4a
Accessible binding site
Inaccessible binding site
b
20,000
15,000
10,000
5,000
0
### Inaccessible genome-wide TFBSs
FoxA1
Hnf4a
Remained closed
Opened
c
75
60
45
30
15
0
### TFBSs around TF-activated genes
FoxA1
Hnf4a
Novel K562 binding site
HepG2 binding site

**Supplemental Fig. 3: Characterization of FoxA1 and Hnf4a binding patterns in FoxA1 or Hnf4a individual clones. (a)** The number of genome-wide FoxA1 or Hnf4a TFBSs that overlap with an ATAC-seq peak in the uninduced (-dox) cells ("Accessible binding site") or that do not overlap with an ATAC-seq peak ("Inaccessible binding site"). **(b)** The number of inaccessible binding sites from (A) that overlap with an ATAC-seq peak in the induced (+dox) cells ("Opened") or that do not overlap with an ATAC-seq peak ("Remained closed"). **(c)** The number of FoxA1 or Hnf4a binding sites within 50 kb of each FoxA1- or Hnf4a-activated gene characterized as either a "HepG2 binding site," where the TFBS overlaps a TFBS of FoxA1 or Hnf4a in HepG2 liver cells, or as a "Novel K562 binding site," where the TFBS does not overlap with a HepG2 binding site.

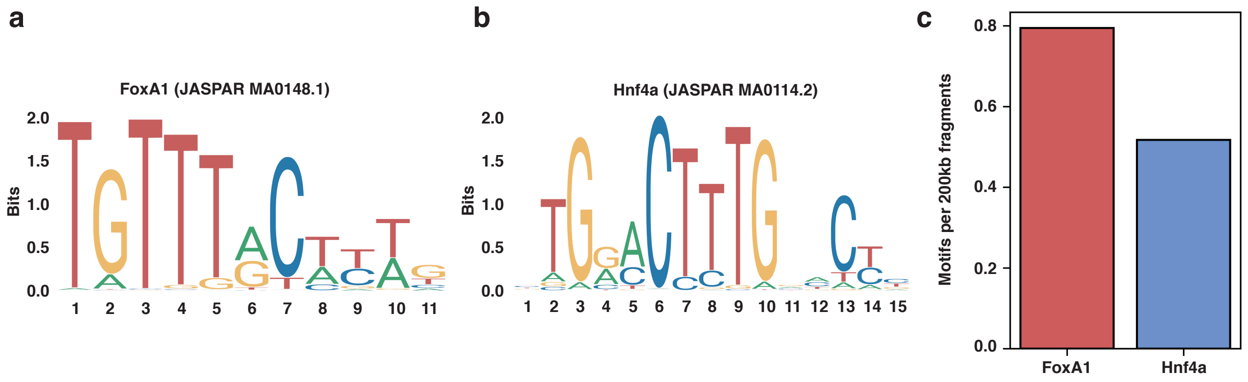
a
FoxA1 (JASPAR MA0148.1)
Bits
2.0
1.5
1.0
0.5
0.0
1 2 3 4 5 6 7 8 9 10 11
b
Hnf4a (JASPAR MA0114.2)
Bits
2.0
1.5
1.0
0.5
0.0
1 2 3 4 5 6 7 8 9 10 11 12 13 14 15
c
Motifs per 200kb fragments
0.8
0.6
0.4
0.2
0.0
FoxA1
Hnf4a

**Supplemental Fig. 4: FoxA1 and Hnf4a motif scanning. (a)** Human FoxA1 sequence logo from JASPAR. **(b)** Human Hnf4a sequence logo from JASPAR. **(c)** 1,000 random 200 bp fragments were generated using Bedtools and then scanned for FoxA1 and Hnf4a motifs with FIMO using 1e-3 a p-value threshold. Total motif count was divided by the number of non-N containing random sequences (924) to identify motifs per random 200bp fragment.

a

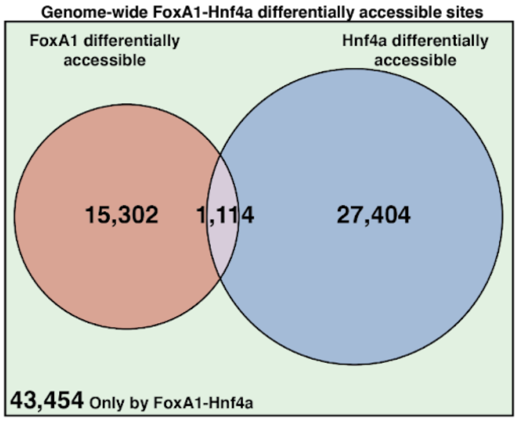
Genome-wide FoxA1-Hnf4a differentially accessible sites
FoxA1 differentially accessible
Hnf4a differentially accessible
15,302
1,114
27,404
43,454 Only by FoxA1-Hnf4a

**Supplemental Fig. 5: Characterization of FoxA1-Hnf4a differential accessibility. (a)** Venn diagram of all FoxA1-Hnf4a induced differentially accessible peaks categorized by whether the peak was also induced in the FoxA1 clone, Hnf4a clone, neither, or both.

#### Supplementary Tables

**Supplementary Table 1**. FoxA1 Gene Ontology Analysis.

| FoxA1 overrepresented biological process | Representative genes | FDR |
|---|---|---|
| Fibrinolysis | SERPINF2, SERPING1, FGB, FGG, F2 | 9.48e-07 |
| Negative regulation of complement activation | SERPINF2, VTN, F2 | 4.36e-02 |
| Regulation of heterotypic cell-cell adhesion | APOA1, FGB, FGG | 6.24e-03 |
| Acute phase response | ITIH4, F2, SERPINF2 | 5.75e-04 |
| Platelet degranulation | ALB, FGB, FGG, SERPINF2 | 4.00e-06 |

**Supplementary Table 2**. Hnf4a Gene Ontology Analysis.

| Hnf4a overrepresented biological process | Representative genes | FDR |
|---|---|---|
| Negative regulation of cholesterol import | APOC3, APOA2 | 5.92e-03 |
| Negative regulation of VLDL particle remodeling/clearance | APOA1, APOC3, APOA2 | 1.97e-04 |
| Chylomicron assembly/remodeling | APOC2, APOC3, APOA1, APOA2 | 2.22e-05 |
| Tyrosine catabolic process | HGD, HPD | 1.61e-02 |
| Phospholipid efflux | APOC2, APOC3, APOA1, APOA2 | 4.14e-05 |

**Supplementary Table 3**. Primer sequences.

| Primer | Sequence |
|---|---|
| foxa1-v5-step1-R | agagggttagggataggcttaccacttgtattcaaaactggtcg |
| hnf4a-v5-step1-R | agagggttagggataggcttaccagcaacttgcccaaagcggc |
| v5-step2-F | ctacgtagaatcgagaccgaggagagggttagggataggctt |
| foxa1-pinducer-F | ccagcctccgcggccccgaaatgttgggcaccgtgaag |
| foxa1-pinducer-R | tgggacgtcgtatgggtattctacgtagaatcgagaccg |
| hnf4a-pinducer-F | ccagcctccgcggccccgaaatgcgactctccaaaacc |
| hnf4a-pinducer-R | tgggacgtcgtatgggtattctacgtagaatcgagacc |
| foxa1-qpcr-F | catgagacaagcgactggaa |
| foxa1-qpcr-R | tattaaaggaggccggtgtc |
| alb-qpcr-F | ctgcctgcctgttgccaaagc |
| alb-qpcr-R | ggcaaggtccgccctgtcatc |
| hnf4a-qpcr-F | aatgacacgtccccatcaga |
| hnf4a-qpcr-R | ggagtacatgtggttcttcc |
| apob-qpcr-F | agaggacagagccttggtggat |
| apob-qpcr-R | ctggacaaggtcatactctgcc |
| hprt-qpcr-F | tgacactggcaaaacaatgca |
| hprt-qpcr-R | ggtccttttcaccagcaagct |

**Supplementary Table 4**. Antibodies.

| **Target** | **Species** | **Type** | **Catalog#** |
|---|---|---|---|
| Human FoxA1 | Rabbit | Monoclonal | Cell Signaling #53528 |
| Human Hnf4a | Mouse | Monoclonal | Invitrogen #MA1-199 |
| Histone H3K4me3 | Rabbit | Polyclonal | Epicypher #13-0041 |
| Rabbit | Goat | Polyclonal | Epicypher #13-0047 |
| Mouse | Goat | Polyclonal | Epicypher #13-0048 |
